# Bovine dendritic-cell subsets in lymphoid and non-lymphoid tissues: work in progress

**DOI:** 10.1101/2024.09.17.613501

**Authors:** SC Talker, GT Barut, A Summerfield

## Abstract

Bovine dendritic-cell subsets can be readily identified in flow cytometry by staining for Flt3, CD4 and CD13. While this gating strategy could be confirmed by bulk RNA-seq of subsets sorted from peripheral blood mononuclear cells, it remains to be determined if the same gating strategy can be applied to gate on cDC1, cDC2 and pDC within lymphoid and non-lymphoid tissues. With this preprint we aim to inform on the current status of phenotyping experiments performed on Flt3^+^ cells in mesenteric lymph nodes, spleen and lung of cattle. Distinct DC subsets can be gated in all tissues based on CD4 and CD13, with high frequencies of Flt3^+^CD13^+^ cells (putative cDC1) detected in the lung. Further phenotyping performed on lymph node cells (in particular CADM1, CD26 and CD123) support the subset classification analogous to blood, with pDC being Flt3^+^CD4^+^CD13^-^ and cDC1 being Flt3^+^CD4^-^CD13^+^. Heterogeneous expression of CADM1 and CD26 on Flt3^+^CD4^-^CD13^-^ putative cDC2 warrants further investigations.

## Introduction

Across species, dendritic cells (DC) can be classified into at least three subsets encompassing conventional dendritic cells of type 1 and 2, and plasmacytoid dendritic cells (Auray et al., 2016; Summerfield et al., 2015; Talker et al., 2018). Given their important role in the initiation and shaping of T-cell responses and in antiviral immunity (Durai and Murphy, 2016), their enumeration and phenotypic characterization by flow cytometry is a valuable readout for example in infection experiments performed on cattle.

A combination of only 3 markers (Flt3, CD4, CD13) was shown to enable detection of cDC1 (Flt3^+^CD4^-^CD13^+^), cDC2 (Flt3^+^CD4^-^CD13^-^), and pDC (Flt3^+^CD4^+^CD13^-^) in blood of cattle (Talker et al., 2018).

Staining for the DC marker Flt3 (fms-like tyrosine kinase 3) is key, although Flt3-independent staining panels have been designed based on extensive phenotyping of above-mentioned subsets in blood of cattle (Barut et al., 2020; Talker et al., 2018). Accordingly, when gating on large cells (FSC-A vs. SSC-A), cDC1 can be gated as CD172a^-/dim^CD13^+^CD26^+^, cDC2 as CD172a^+^CD14^-^CD16^-^, and pDC as CD11c^-^CD4^+^CD5^+^.

To which extent these phenotypic definitions are valid in tissues is currently unknown. The phenotyping experiments we performed indicate that Flt3 staining is possible in lymph node and spleen, and that CD4/CD13-defined subsets likely correspond to the DC subsets described in blood. However, staining of non-lymphoid tissues that were digested using collagenase, such as the lung, may require additional strategies to enhance the signal of Flt3 staining.

## Materials and Methods

### Isolation of peripheral blood mononuclear cells (PBMC)

Blood collection and PBMC isolation were performed as previously described (Talker et al., 2018). The blood sampling was performed in compliance with the Swiss animal protection law and approved by the animal welfare committee of the Canton of Bern, Switzerland, license number BE102/15.

### Cell isolation from lymph nodes and spleen

Mesenteric lymph nodes and spleen were collected into ice-cold PBS containing 1 mM EDTA (Invitrogen) (PBS/EDTA). Tissue was cut into small pieces and minced using GentleMACS C tubes (Miltenyi Biotec Swiss AG, Solothurn, Switzerland). Minced tissue was passed through a sieve and centrifuged (300 x g, 10 min, 4°C). This was followed by an incubation in 0.1 mg/mL DNase I (Worthington-Biochem, BioConcept, Basel, Switzerland) solution, for 15 min at room temperature and subsequent passage through a cell strainer (70 μm). After another washing step (300 x g, 10 min, 4°C), the cell pellet was resuspended in PBS/EDTA and was layered onto Ficoll Paque (1.077 g/mL; GE Healthcare Europe GmbH) and centrifuged (800 x g, 25 min) at room temperature. Finally, cells were washed twice with cold PBS/EDTA (400 x g, 10 min, 4°C), and counted.

### Cell isolation from lungs

Parts of lungs were collected into ice-cold PBS containing 1 mM EDTA (Invitrogen) (PBS/EDTA). In order to remove blood, lung pieces were washed with cold PBS/EDTA several times. Tissue was cut into small pieces and digested using an enzyme solution [1.25 mg/mL Collagenase I (Worthington-Biochem, BioConcept, Basel, Switzerland), 2.5 mg/mL Collagenase II (Worthington-Biochem), 1.6 mg/mL DNase I (Worthington-Biochem) in DMEM supplemented with 5% FBS (Gibco)] by incubating at 37°C for 2 hours on a shaker. This was followed by a mincing step using GentleMACS C tubes (Miltenyi Biotec Swiss AG, Solothurn, Switzerland), and density gradient centrifugation using Ficoll as described above for lymph nodes and spleen.

### MACS and flow cytometry

Isolated cells were depleted of CD3^+^ cells by magnetic sorting (Miltenyi Biotec) to enable a clear identification of pDC that express low levels of Flt3 and share common surface markers with CD4^+^ T cells.

Six-color staining was performed in 96-well U-bottom plates with 1 × 10^7^ cells per sample and encompassed five incubation steps, each for 20 min at 4°C. Washing steps between incubations were done with Cell Wash (BD Biosciences, Allschwil, Switzerland).

Antibodies and reagents are listed in Table 1. Bovine recombinant IL-3 was produced in house (unpublished; M&M provided in updated version of the preprint).

**Table 1.**
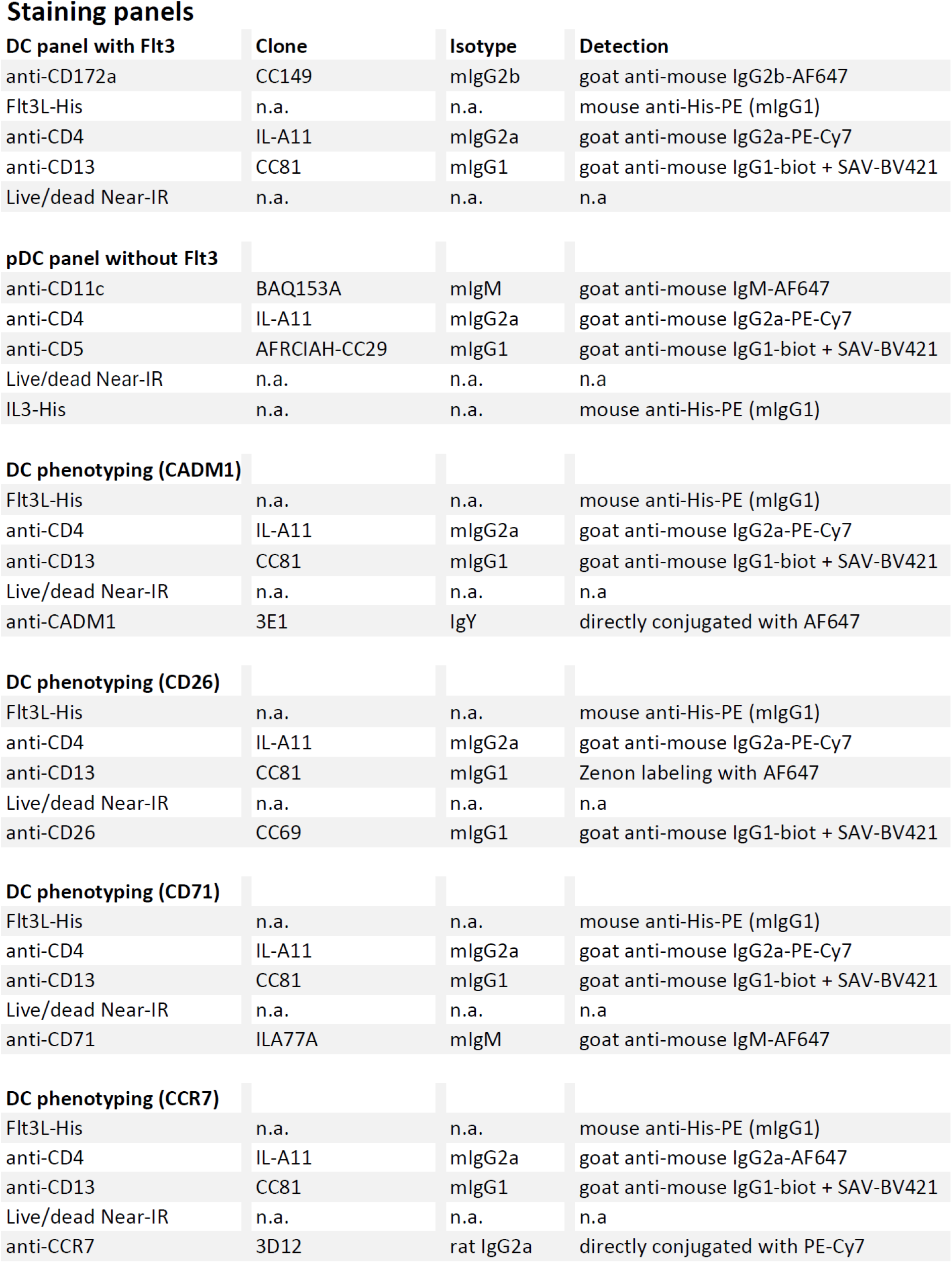

**Staining panels**
| **DC panel with Flt3** | **Clone** | **Isotype** | **Detection** |
| --- | --- | --- | --- |
| anti-CD172a | CC149 | mlgG2b | goat anti-mouse IgG2b-AF647 |
| Flt3L-His | n.a. | n.a. | mouse anti-His-PE (mlgG1) |
| anti-CD4 | IL-A11 | mlgG2a | goat anti-mouse IgG2a-PE-Cy7 |
| anti-CD13 | CC81 | mlgG1 | goat anti-mouse IgG1-biot + SAV-BV421 |
| Live/dead Near-IR | n.a. | n.a. | n.a |
| **pDC panel without Flt3** |  |  |  |
| anti-CD11c | BAQ153A | mlgM | goat anti-mouse IgM-AF647 |
| anti-CD4 | IL-A11 | mlgG2a | goat anti-mouse IgG2a-PE-Cy7 |
| anti-CD5 | AFRCIAH-CC29 | mlgG1 | goat anti-mouse IgG1-biot + SAV-BV421 |
| Live/dead Near-IR | n.a. | n.a. | n.a |
| IL3-His | n.a. | n.a. | mouse anti-His-PE (mlgG1) |
| **DC phenotyping (CADM1)** |  |  |  |
| Flt3L-His | n.a. | n.a. | mouse anti-His-PE (mlgG1) |
| anti-CD4 | IL-A11 | mlgG2a | goat anti-mouse IgG2a-PE-Cy7 |
| anti-CD13 | CC81 | mlgG1 | goat anti-mouse IgG1-biot + SAV-BV421 |
| Live/dead Near-IR | n.a. | n.a. | n.a |
| anti-CADM1 | 3E1 | IgY | directly conjugated with AF647 |
| **DC phenotyping (CD26)** |  |  |  |
| Flt3L-His | n.a. | n.a. | mouse anti-His-PE (mlgG1) |
| anti-CD4 | IL-A11 | mlgG2a | goat anti-mouse IgG2a-PE-Cy7 |
| anti-CD13 | CC81 | mlgG1 | Zenon labeling with AF647 |
| Live/dead Near-IR | n.a. | n.a. | n.a |
| anti-CD26 | CC69 | mlgG1 | goat anti-mouse IgG1-biot + SAV-BV421 |
| **DC phenotyping (CD71)** |  |  |  |
| Flt3L-His | n.a. | n.a. | mouse anti-His-PE (mlgG1) |
| anti-CD4 | IL-A11 | mlgG2a | goat anti-mouse IgG2a-PE-Cy7 |
| anti-CD13 | CC81 | mlgG1 | goat anti-mouse IgG1-biot + SAV-BV421 |
| Live/dead Near-IR | n.a. | n.a. | n.a |
| anti-CD71 | ILA77A | mlgM | goat anti-mouse IgM-AF647 |
| **DC phenotyping (CCR7)** |  |  |  |
| Flt3L-His | n.a. | n.a. | mouse anti-His-PE (mlgG1) |
| anti-CD4 | IL-A11 | mlgG2a | goat anti-mouse IgG2a-AF647 |
| anti-CD13 | CC81 | mlgG1 | goat anti-mouse IgG1-biot + SAV-BV421 |
| Live/dead Near-IR | n.a. | n.a. | n.a |
| anti-CCR7 | 3D12 | rat IgG2a | directly conjugated with PE-Cy7 |

Prior to incubation with primary antibodies, cells were incubated with purified bovine IgG (Bethyl laboratories, Montgomery, USA) in order to block Fc receptors. ChromPure mouse IgG (Jackson ImmunoResearch, Lubioscience GMBH, Zürich, Switzerland) was used in the fourth step to block remaining binding sites of isotype-specific secondary antibodies. In the final step, anti-His-PE (Miltenyi Biotec Swiss AG, Solothurn, Switzerland) was added in order to stain His-tagged Flt3L. Bovine Flt3L (NCBI NM_181030.2) was produced as previously described (Ziegler et al., 2016). Samples were acquired with a FACSCanto II flow cytometer (BD Biosciences) equipped with three lasers (405, 488, and 633 nm). Whenever possible, at least 1 × 10^6^ cells were recorded in the “large-cell” gate. Compensation was calculated by FACSDiva software following the measurement of single-stained samples. Highly autofluorescent cells were gated out using the AF488 channel (empty) vs. the PE channel (Figure 1).

**Figure 1.**
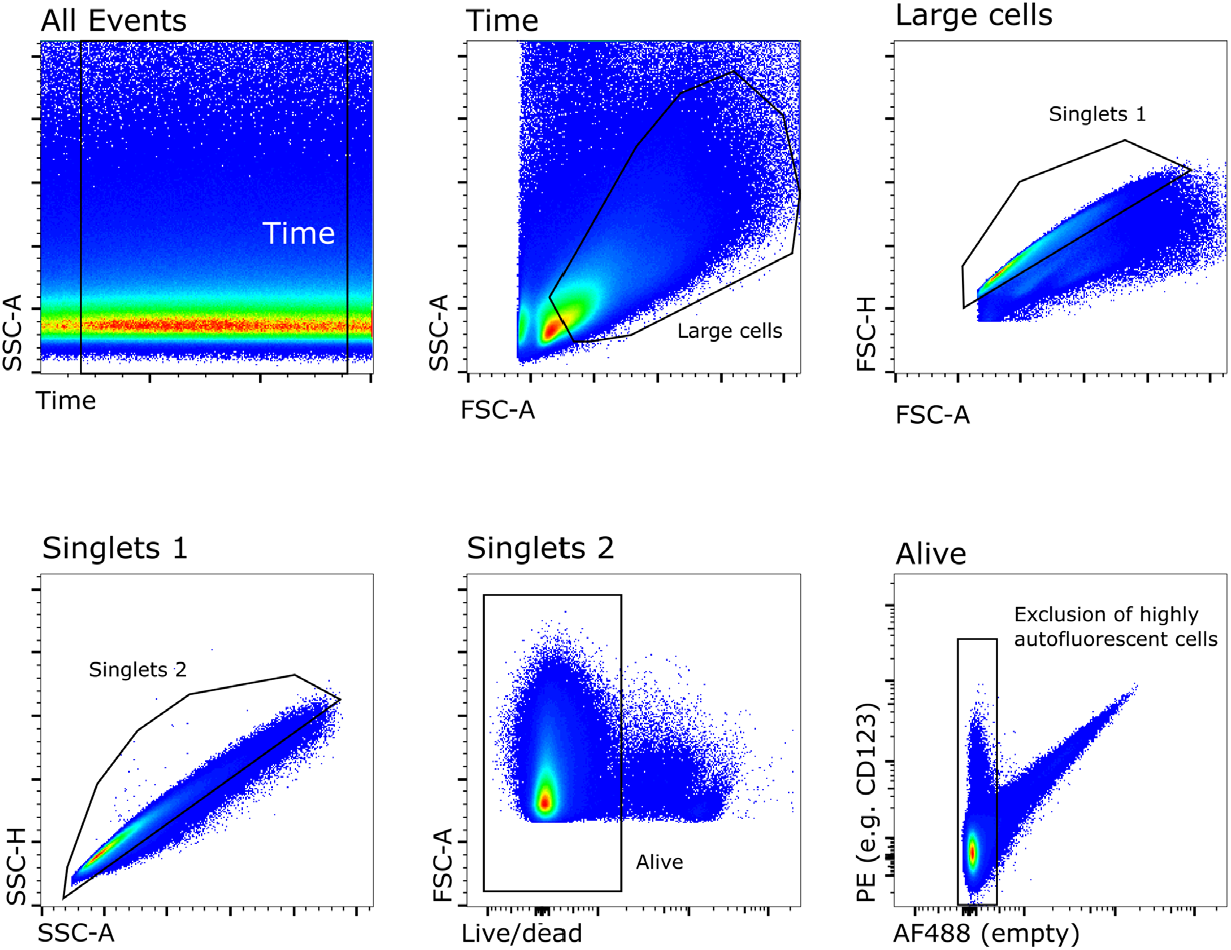
Gating strategy and exclusion of highly autofluorescent cells. Following the exclusion of irregular events using the “Time” parameter, large mononuclear cells were gated based on FSC-A and SSC-A. Single cells were gated in the FSC-A vs. FSC-H plot (Singlets 1) and the SSC-A vs. SSC-H plot (Singlets 2). Viable cells were gated based on weak staining with live/dead stain. Within viable cells, highly autofluorescent cells were excluded based on a AF488 vs. PE plot, with AF488 used as an empty channel.

## Results & Discussion

Recombinant bovine Flt3L enables detection of bovine dendritic cells in blood, which can be further differentiated based on CD4 and CD13 expression (Talker et al., 2018).

Phenotyping experiments reported here indicate that this three-marker panel (Flt3, CD4, CD13) is likely suited to identify corresponding subsets in lymph node, spleen and lung.

Notably, when staining cells isolated from tissues, depletion of CD3^+^ cells is beneficial, in particular for gating on pDC who express low levels of Flt3. We previously reported on a Flt3-independent staining panel for pDC, including CD11c and CD5 alongside CD4, where bovine pDC in blood can be gated as CD11c^-^CD5^+^CD4^+^ within CD3-depleted cells (Barut et al., 2020). Our results indicate that this strategy can also be applied for pDC from bovine lymph nodes (Figure 2).

**Figure 2.**
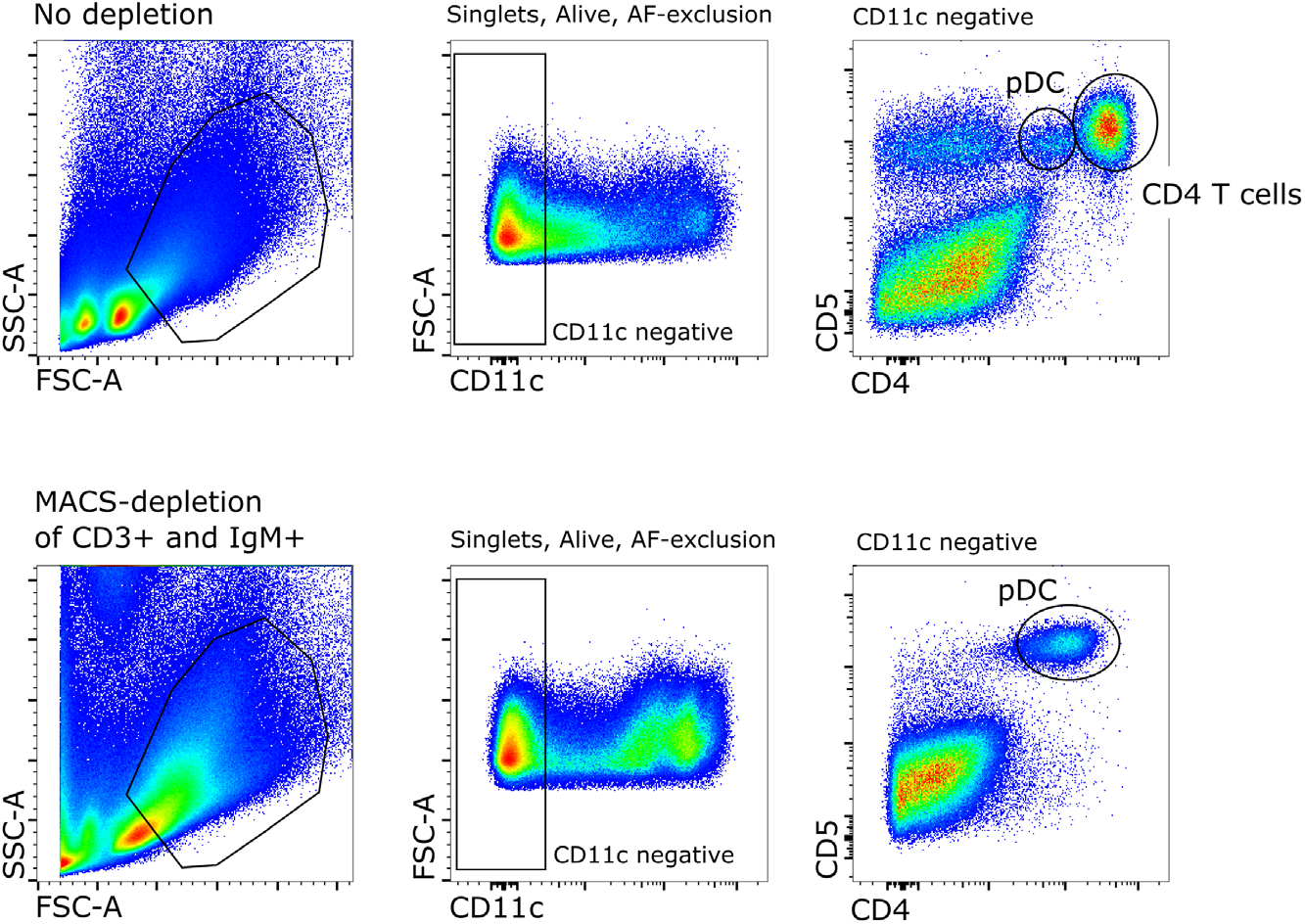
Depletion of CD3^**+**^ cells and Flt3-independent gating of pDC in lymph nodes. Flt3-independent gating is shown prior to depletion (upper panel) and following depletion of CD3^+^ and IgM^+^ cells by MACS (lower panel). Gating was performed as shown in Figure 1. Putative pDC were gated as CD4^+^CD5^+^ cells within the CD11c negative gate.

Staining of bovine lymph node and spleen with Flt3L-His followed by anti-His-PE (mouse IgG1) yielded acceptable Flt3 detection (Figure 3). However, staining of DC from lung tissue requires further optimization. Likely, addition of anti-mouse IgG1-PE will significantly improve the Flt3 staining, as observed with experiments performed on caprine lymph nodes (unpublished). Backgating shown in Figure 3 illustrates that CD4/CD13 defined subsets form uniform populations that localize as expected in the CD172a vs. Flt3 plot: highest Flt3 expression on putative cDC1, lowest Flt3 expression on putative pDC, and highest CD172a expression with intermediate Flt3 expression on putative cDC2.

**Figure 3.**
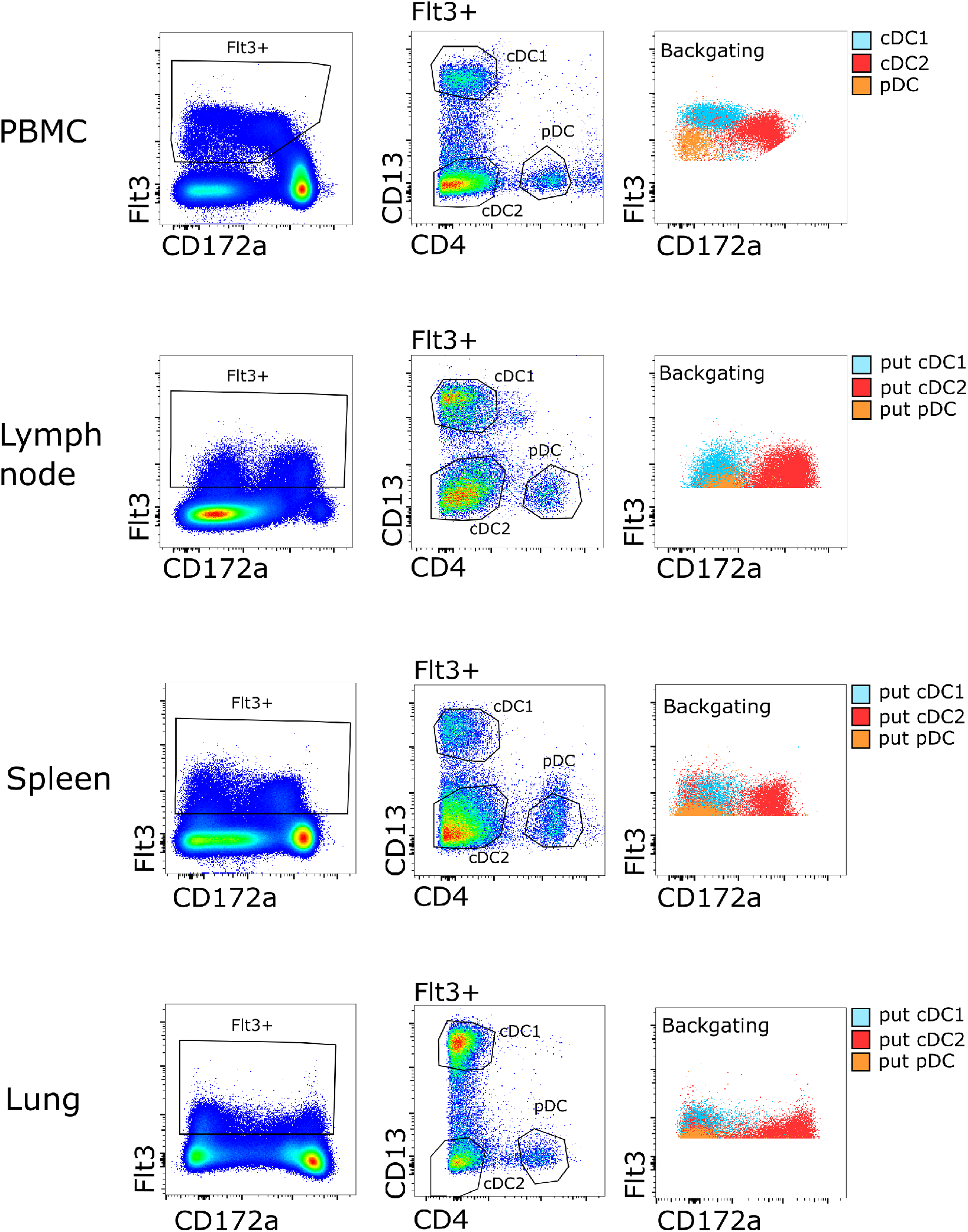
Gating of DC subsets in blood, lymph node, spleen, and lung. Freshly isolated cells were stained for flow cytometry and pre-gated as shown in Figure 1. Based on the gating strategy established for bovine DC subsets in blood (PBMC), putative DC subsets in lymph node, spleen and lung were gated as Flt3^+^CD4^+^CD13^-^ (pDC), Flt3^+^CD4^-^CD13^+^ (cDC1) and Flt3^+^CD4^-^CD13^-^ (cDC2). Backgating indicates location of subsets on the CD172a vs. Flt3 plot.

Further phenotyping of DC subsets derived from bovine lymph nodes, supports this classification. As reported for cDC1 in bovine blood (Talker et al., 2018), putative cDC1 in bovine lymph nodes expressed the highest levels of CADM1 and CD26 (Figure 4A). Moreover, putative CD11c^-^CD4^+^CD5^+^ pDC exclusively expressed CD123 (Figure 4B).

**Figure 4.**
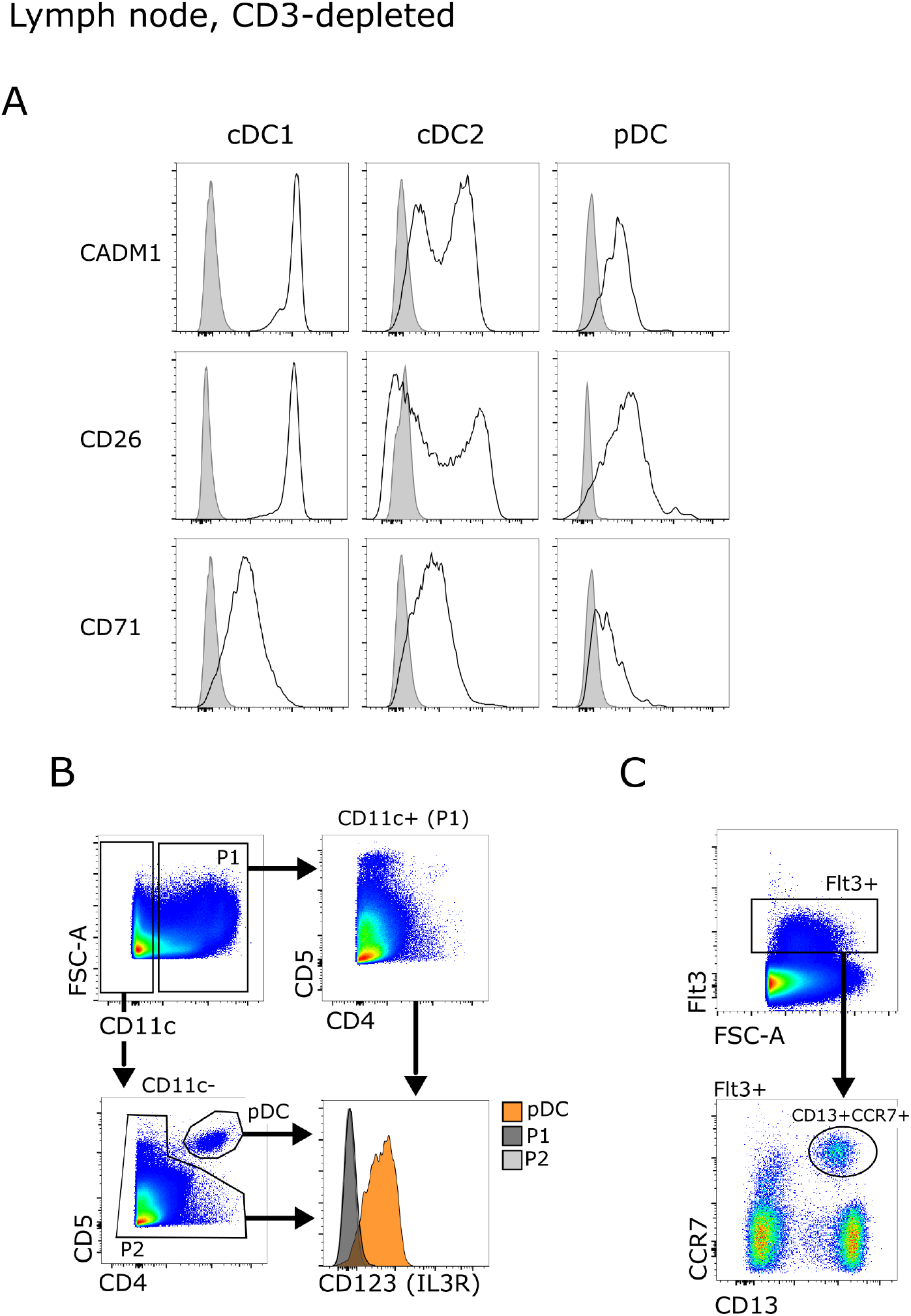
Phenotyping of DC subsets isolated from bovine lymph nodes. Freshly isolated and CD3-depleted cells from bovine mesenteric lymph nodes were stained for flow cytometry and pre-gated as shown in Figures 1 and 3. (A) Expression of CADM1, CD26, and CD71 on putative cDC1, cDC2, and pDC. Gray histograms represent FMO controls. (B) Expression of CD123 on pDC gated as CD11c^-^CD4^+^CD5^+^ (orange histogram). Overlaying gray histograms show CD123 expression on CD11c^+^ cells (P1), and on non-pDC within CD11c^-^ cells (P2). (C) Expression of CCR7 on Flt3^+^CD13^+^ putative cDC1. Data are representative for at least two animals.

Chemokine receptor 7 (CCR7) was previously shown to be upregulated on blood DC subsets upon *in vitro* stimulation with TLR ligands, both on protein level and RNA level (Barut et al., 2020). However, when analyzing single-cell RNA-seq data of enriched mononuclear phagocytes from bovine lymph nodes, we could not detect ANPEP (CD13) mRNA in the cluster of CCR7 positive cells (Barut et al., 2023). Phenotyping experiments shown here indicate that CD13 is expressed on CCR7^+^ cells, but is slightly downregulated when compared to CD13^+^CCR7^-^ cells (Figure 4C).

Taken together, our data so far indicate that staining for Flt3, CD4 and CD13 can be used to identify DC subsets in lymph nodes, spleen and lung. Additional phenotyping is necessary to corroborate this gating strategy and to establish Flt3-independent panels in different tissues, including skin and intestine of cattle.

Given the complexity of the DC compartment, including newly defined DC3 and transitional dendritic cells (tDC), the CD4/CD13-based gating strategy can only serve as a rough approximation. This limitation needs to be taken into consideration, especially when performing functional assays on those subsets. However, more fine-grained classification of DC by flow cytometry is very challenging, especially for the cDC2 compartment. Single-cell RNA-seq data on mononuclear phagocytes enriched from blood and tissues of cattle will be highly valuable to estimate the consequences of this “oversimplification”.

Certainly, phenotyping experiments performed on tissue DC will inform the design of enrichment strategies that can be harnessed to zoom into certain compartments by high-dimensional approaches, such as RNA sequencing at the single-cell level.

